## Supplementary material for "Cryo-electron tomography reveals the microtubule-bound form of inactive LRRK2": Data Availability

--------------------
GENERAL INFORMATION 
--------------------
This readme file was generated on [2024-08-06] by [Siyu Chen]

Title of Dataset: Models and maps of LRRK2I2020T/LRRK2WT with MLi-2/GZD-824 on pre-formed microtubules
Description of Dataset: Models and maps of LRRK2I2020T with MLi-2 on pre-formed microtubules
Principal Investigator: Siyu Chen,, 0000-0003-4565-4772
Date of Data Collection: 2023-08-12
Software Dependencies: Chimerax

-------------
FILE OVERVIEW 
-------------
Directory of Files: 9CHO, EMD-45591, EMD-45592, EMD-45593, EMD-45594, EMD-45595, EMD-45596
Relationship Between Files: 9CHO is the atomic model generated from EMD maps
File Formats: .cif, .mrc
File Naming Convention: all naming follows the online databank naming convention

-----------
METHODOLOGY
-----------
Description of methods used for data collection: Electron images are collected from transmission electron microscope and subtomograms are picked and averaged to generate the EM maps. The atomic model is generated from homologs structure and dynamic fitting. 

Description of methods used for data processing: The following softwares were used to generate subtomograms from election microscopy images: serialEM, Warp, IMOD, Dynamo in Matlab (with shared home-brew scripts). The following softwares were used to obtain EMD-maps and generate atomic structural models: Relion, ChimeraX, ISOLDE

-----------------------
DATA ACCESS AND SHARING
-----------------------
Publications based on this dataset: Cryo-electron tomography reveals the microtubule-bound form of inactive LRRK2
Recommended citation for this dataset: Use the PDB and EMDB code as indicated
License information: CC BY 4.0.
